# CNSA: a data repository for archiving omics data

**DOI:** 10.1101/2020.04.07.030833

**Authors:** Xueqin Guo, Fengzhen Chen, Fei Gao, Ling Li, Ke Liu, Lijin You, Cong Hua, Fan Yang, Wanliang Liu, Chunhua Peng, Lina Wang, Xiaoxia Yang, Feiyu Zhou, Jiawei Tong, Jia Cai, Zhiyong Li, Bo Wan, Lei Zhang, Tao Yang, Minwen Zhang, Linlin Yang, Yawen Yang, Wenjun Zeng, Bo Wang, Xiaofeng Wei, Xun Xu

**Affiliations:** China National GeneBank, Shenzhen 518120, China; BGI-Shenzhen, Shenzhen 518083, China; Guangdong Provincial Key Laboratory of Genome Read and Write, Shenzhen 518120, China

## Abstract

With the application and development of high-throughput sequencing technology in life and health sciences, massive multi-dimensional biological data brings the problem of efficient management and utilization. Database development and biocuration are the prerequisites for the reuse of these big data. Here, relying on China National GeneBank (CNGB), we present CNGB Sequence Archive (CNSA) for archiving omics data, including raw sequencing data and its analytical data and related metadata which are organized into six objects, namely Project, Sample, Experiment, Run, Assembly, and Variation at present. Moreover, CNSA has created the correlation model of living samples, sample information, and analytical data on some projects, so that all data can be traced throughout the life cycle from the living sample to the sample information to the analytical data. Complying with the data standards commonly used in the life sciences, CNSA is committed to building a comprehensive and curated data repository for the storage, management and sharing of omics data, improving the data standards, and providing free access to open data resources for worldwide scientific communities to support academic research and the bio-industry. Database URL: https://db.cngb.org/cnsa/

## Introduction

In data-intensive science era, life science research is seen as a data-driven, exploration-centered style of science. With the development of sequencing technology, the rapid increase of sequencing throughput and the dramatic drop of per-base cost of raw sequence have made large-scale population genomics research, precision medicine research, and biodiversity research possible, e.g., the UK’s 100,000 Genomes Project (1), the International Cancer Genome Consortium (ICGC) (2), the Cancer Genome Atlas (TCGA) (3), the China Kadoorie Biobank (CKB: https://www.ckbiobank.org/site/) (4) and Earth BioGenome Project (EBP) (5). However, it poses great challenges in big data deposition, integration and sharing, where IT infrastructure and software tools are heavily used.

Many organizations in the world have made great efforts for archiving and sharing of omics data. The International Nucleotide Sequence Database Collaboration (INSDC) (6) represents one of the most celebrated global initiatives in data and associated metadata sharing, which operates between DNA Data Bank of Japan (DDBJ) (7), the European Molecular Biology Laboratory-European Bioinformatics Institute (EMBL-EBI) (8), and the National Center for Biotechnology Information (NCBI) (9). In order to facilitate exchange of information on genomic samples and their derived data, the Global Genome Biodiversity Network (GGBN) Data Standard (10) is intended to provide a platform to promote the efficient sharing and usage of genomic sample material and associated specimen information in a consistent way. Global Alliance for Genomics and Health (GA4GH) (11), an international, nonprofit alliance, already brings together more than 500 leading organizations to accelerate progress in genomic research and human health. In addition, DataCite (12) has developed tools and methods that make data more accessible and more useful. In China, many scientific institutions have also made great efforts and established multiple omics database systems such as the National Genomics Data Center (NGDC: https://bigd.big.ac.cn/), BIGBIM (https://www.biosino.org/bigbim/index), the National Center for Protein Science •Shanghai (NCPSS: http://www.sibcb-ncpss.org/index.action), etc.

The CNGB (13) (https://www.cngb.org/home.html) based in Shenzhen was established in January 2011, which is committed to supporting public welfare, life science research, innovation and industry incubation, through effective bioresource conservation, digitalization and utilization. Based on this concept and relying on the CNGB, China National GeneBank DataBase (CNGBdb: https://db.cngb.org/) is built as a unified platform built for biological big data sharing and application services to the research community. Based on the big data and cloud computing technologies, it provides data services such as archive, analysis, knowledge search, management authorization, and visualization. CNSA is the data archiving system of CNGBdb and is built for archiving omics data including not only raw sequencing data and its analysis result data, but also related metadata. At present, CNSA follows the data standards and structures of INSDC, DataCite, GA4GH and GGBN for data compatibility and provides global users with data archival and sharing services of omics data such as data submission, data storage, data retrieval and data reference. All archived public data is freely available to worldwide scientific communities.

## Methods

### System structure

Based on the Django framework (Django is a high-level Python Web framework that encourages rapid development and clean, pragmatic design, https://www.djangoproject.com/), CNSA is developed in Python. In order to provide more stable and fast services, CNSA server is built on the Centos-7 operating system with the following six servers: NGINX for providing static resource access, uWSGI for deploying services, PostgreSQL and MongoDB for supporting metadata storage, FTP server for uploading and storing data files and Elasticsearch for data retrieval. In addition, Redis-based caching system is used to help improve data verification speed.

### Data security

Relying on the data room of CNGB and CNGBdb that have passed the three-level review of information security level protection and the protection capability review of trusted cloud service, CNSA adopts corresponding security technologies in user access, data room, firewall, application architecture, database, and data storage. The entire site of the CNSA uses https to encrypt user access requests to prevent stealing and tampering during data transmission. Django ORM is used to avoid SQL injection, and fields retrieved from the database are filtered before being displayed to prevent XSS attacks. All information submitted by users is submitted in Post mode, and Django’s CsrfViewMiddleware is used to prevent CSRF attacks. The firewall security technology of the CNGB data room can ensure the legality of data access. Moreover, CNSA adopts high-performance distributed object storage for data archiving, and combines a high-availability and disaster tolerance backup storage system to ensure data storage security, and the database is backed up daily and can be restored in a timely manner.

## Results

### Data objects and structure

At present, CNSA follows the data standards and structures of INSDC, DataCite, GA4GH and GGBN to ensure data compatibility, and all data are organized into six objects, i.e., Project, Sample, Experiment, Run, Assembly and Variation. The definitions of data objects and main fields used to describe the data objects are listed in Table 1. As is illustrated in Figure 1, project and sample can be submitted independently, a projects can also be associated with one or more projects, and a sample can also be associated with one or more of experiments, assemblies, or variations. An experiment can be associated with one or more runs. Moreover, each data is assigned a corresponding accession number that can be referenced. Also note that projects and samples are not directly related, and they are not related until experiments, assemblies, or variations are associated with projects and samples, respectively.

**Table 1.** Definitions and main fields of data objects.

| Data object | Definition | Main fields |
| --- | --- | --- |
| Project | An overall description of a single research initiative | project name, project title, public description, sample scope, data type, submitter, funding information, publication |
| Sample | A description of biological source material | sample type, sample name, organism, taxonomy ID, collection |
| Experiment | A description of sample-specific sequencing library, instrument and sequencing methods | file type, sequencing platform, library strategy |
| Run | A description of the sequencing data files that belong to the related experiment | file name, MD5 value |
| Assembly | A collection of genomic sequences that are used to represent the genome of an organism. | molecule type, coverage, sequencing technology, assembly method |
| Variation | Genome variations of any species | variation type, position, variation, detection method, clinical significance, phenotype |

**Figure 1.**
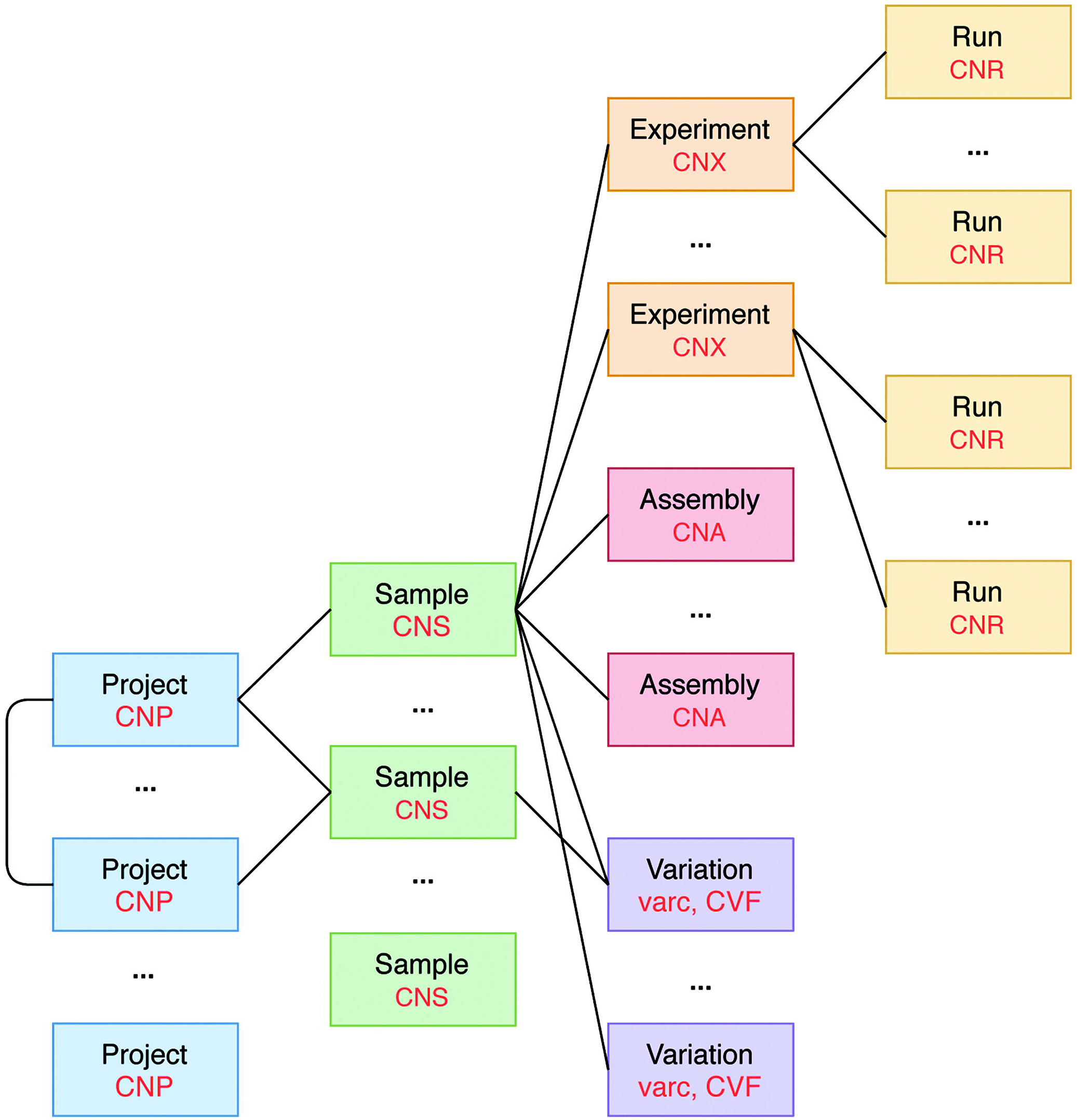
Data model in CNSA.

At present, CNSA has six data objects, and the corresponding prefixes of accession numbers are marked in red.

It’s worth mentioning that CNSA has created the correlation model of living samples, sample, and analytical data on some projects such as the Ruili Botanical Garden project (https://db.cngb.org/search/project/CNPhis0000538) (14) and Culturable Genome Reference (https://db.cngb.org/search/project/CNP0000126/) (15). Figure 2 illustrates the interrelationship of the living samples, sample, and analytical data for the Ruili Botanical Garden project. CNSebb is the prefix of the accession number of the living samples which can be applied to the E-BioBank (EBB: https://db.cngb.org/ebb/), a shared platform for sample resources in the CNGB. The living samples and the sample information are correlative, and the sample information and the analytical data are also correlative, so that all data can be traced throughout the life cycle from the living sample to the sample information to the analytical data.

**Figure 2.**
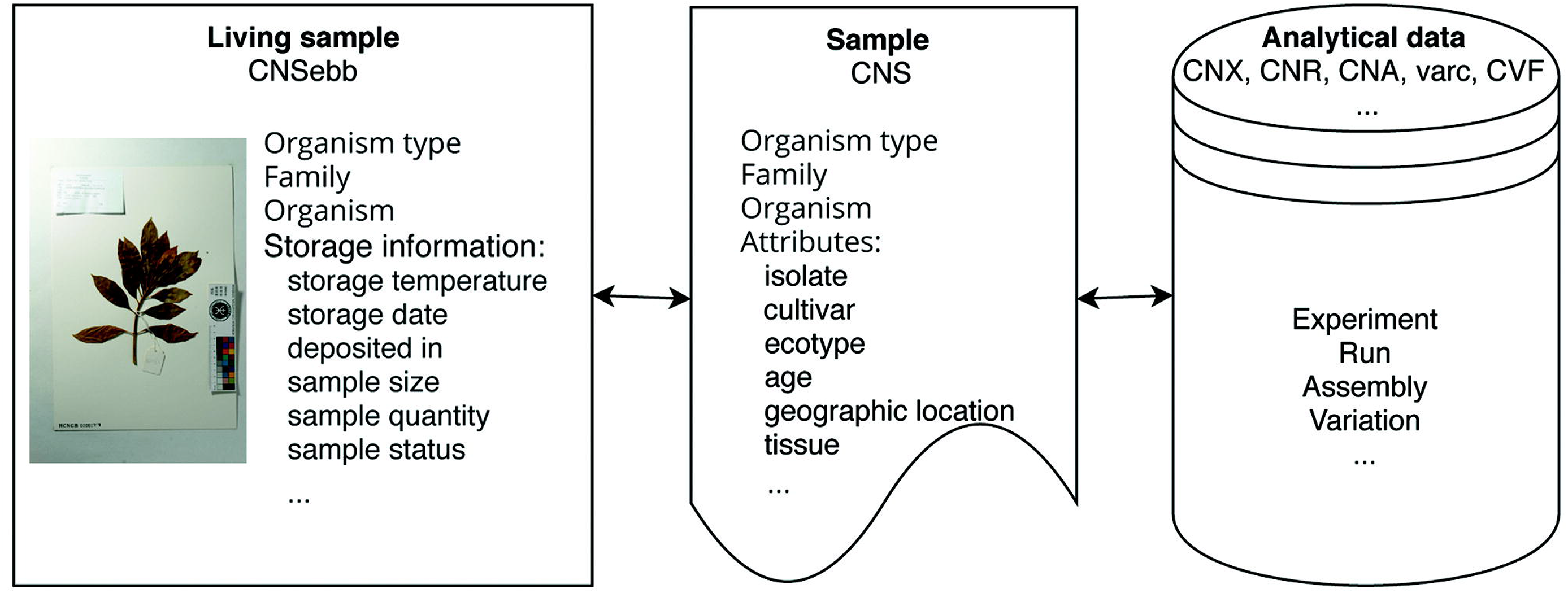
Correlation model for Ruili Botanical Garden project.

### Data submission and curation

Users need to register, login and fill in the submitter’s information before creating a submission. Generally, the order of data submission is project, sample, and related experiments, runs, assemblies, and variations (Figure 3). The data to be submitted includes metadata and data files. At its most basic level, metadata is “data about data”. Here metadata is data that describes a data object, such as the attributes listed in the “Main Fields” column of Table 1. To collect metadata, CNSA provides a user-friendly and easy-to-use submission process that supports Chinese and English bilingual interfaces. It is worth mentioning that CNSA supports batch submission of samples, experiments, runs, assemblies and variations. Therefore, users can fill in a batch submission template for data and submit a batch of data in one submission process. Compared to a single submission, batch submission greatly improves the efficiency of data submission. Users can choose single submission or batch submission according to the number of entries. To simplify the submission of data files, CNSA supports data files to be uploaded via FTP. Moreover, in order to ensure the integrity of the submitted data, users need to submit the corresponding MD5 checksums of data files while submitting the metadata. The system will also automatically check the standardization of some field information submitted by users, such as format, character limit, etc., and verify the MD5 checksums of the data files, and give corresponding prompts if there are errors. After the data is submitted successfully, each piece of data will be assigned an accession number, and each project will be assigned a DOI which is a persistent identifier.

**Figure 3.**
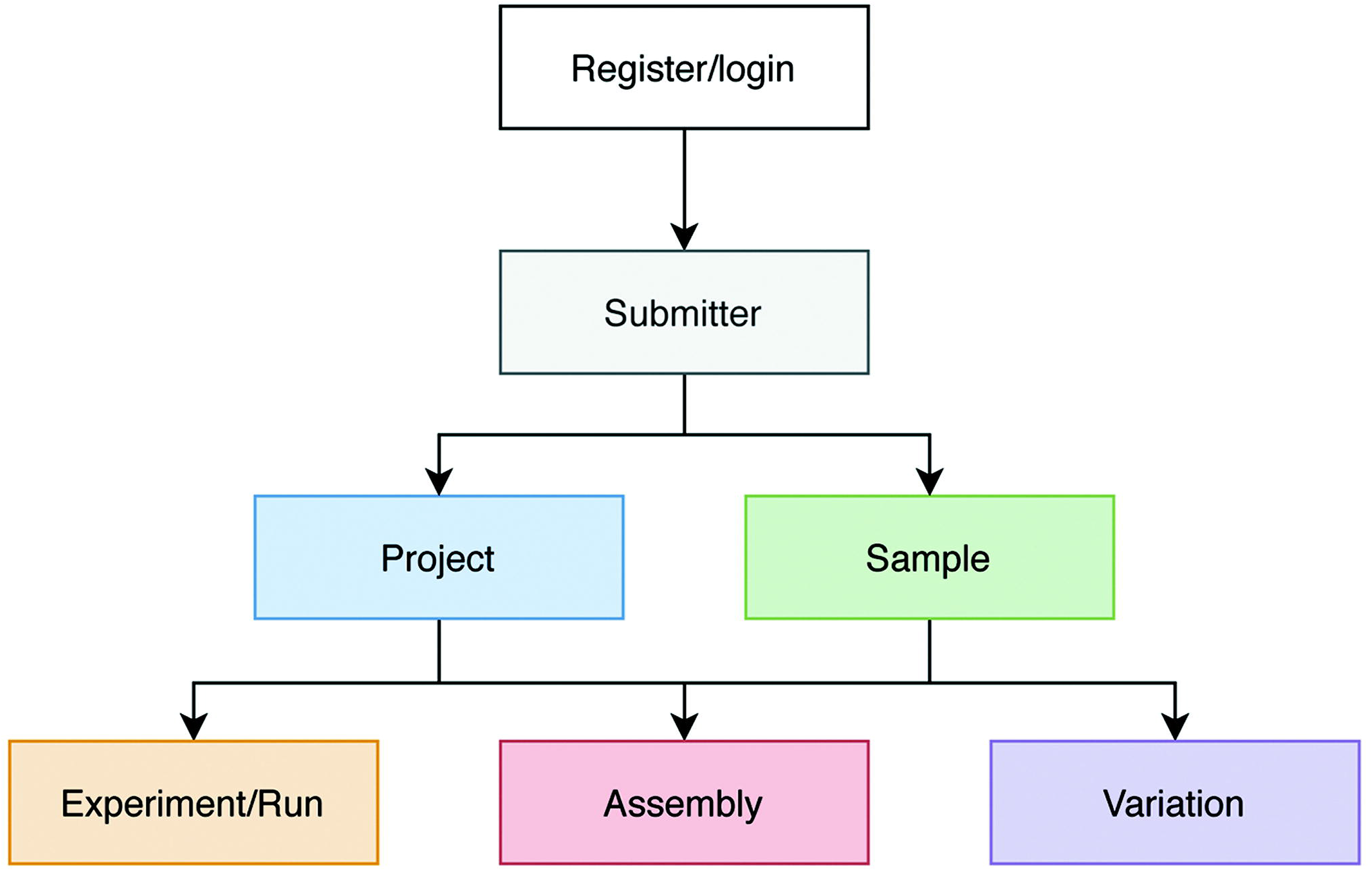
Process of data submission to CNSA.

In addition, like INSDC members, there are two data management manners for projects, public and controlled. The data submitter can choose a data management manner of the project. All metadata and data files associated with the project set to be public will be public on the release date set by the submitter. The public data will be open to the world, and users can access or use it freely without logging in or registering. However, the metadata associated with the project set to be controlled will be public on the metadata release date set by the submitter, and the data files will be under controlled access. Other registered users can submit an application to the CNGB Data Access (CDA: https://db.cngb.org/data_access/) to apply for access to controlled data. Data applicants can use the controlled data only after the data access application has been reviewed and approved. After the project is successfully submitted, it will undergo legality and compliance review such as ethics review and human genetic resources review. All submitted data must pass the legality and compliance review before data biocuration and can only be archived and released or controlled after it has been approved by biocurators. In addition to online check of standardization of some field information, the metadata will also be manually reviewed to ensure its completeness, relevance, and correctness. In order to increase the reusability of data, CNSA only accepts data files in commonly used formats, such as FASTQ, BAM, FASTA, VCF. Moreover, quality control such as checking the correctness of the formats and statistic the data quality using fastp (16) is performed on raw sequencing data files in FASTQ format.

### Data archive and statistics

Currently, CNSA archives biological data and its metadata from around the world, including six objects (Project, Sample, Experiment, Run, Assembly, Variation). The data is made public or controlled based on the submitter’s settings. To ensure data security, CNSA adopts high-performance distributed object storage for data archiving, and double data backup on physically independent disks.

As of March, 2020, CNSA has archived a total of 2073 projects, 308,995 samples, 206,205 experiments, 327,300 runs, and 13,567 assemblies for 2950 species (Figure 4A), submitted by 470 submitters from 101 institutions. The total amount of archived run files and assembly files have reached 2460 TB (Figure 4B). Moreover, CNSA has supported 115 articles published in 79 magazines.

**Figure 4.**
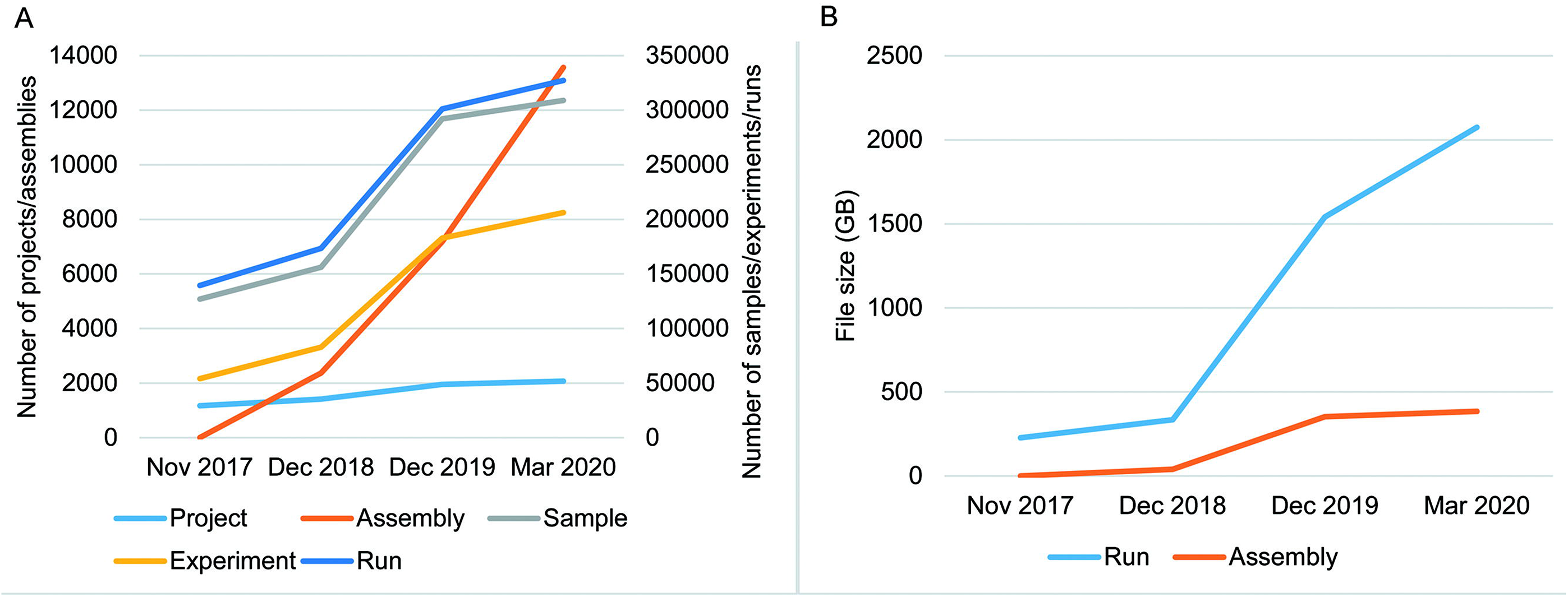
Data statistics of CNSA. **A.** Numbers of Projects, Samples, Assemblies, Experiments and runs in CNSA. **B.** File sizes of Runs and Assemblies in CNSA. All statistics are based on data submitted from November 2017 to March 2020.

### Data retrieval and reference

As mentioned in the submission process, each piece of data will be assigned an accession number, and each project will be assigned a DOI which is a persistent identifier. All public data can be searched in CNSA by data accession numbers of Project, Sample, Experiment, Run, Assembly or any other combination of keywords, and all public data files can be freely accessed through the FTP site (ftp://ftp.cngb.org/). Moreover, both the data accession number and DOI assigned by CNSA can be used to reference the submitted data to support the publication of scientific research results.

## Conclusions and Perspectives

In conclusion, CNSA is a data repository for archiving omics data, including raw sequencing data and its analysis result data and related metadata. Currently, online submissions of projects, samples, experiments, runs, assemblies and variations are accepted. Moreover, compared with similar databases, an advantage worth mentioning is that CNSA has created the correlation model of living samples, sample information, and analytical data on some projects. From now on, CNSA will practice the correlation model on more projects to make all data can be traced throughout the life cycle from the living sample to the sample information to the analytical data and promote the scientific and rational use of biological living samples. All public data resources of CNSA are freely worldwide scientific communities. In compliance with data standards commonly used in the life sciences, CNSA is committed to building a comprehensive and curated data repository for the storage, management and sharing of omics data, and improving the data standards to alleviate the growing management pressure of biological big data and support academic research and the bio-industry.

In order to promote the sharing and exchange of information, technology, and resources of life science under the unified standards and regulations that are formed, and facilitate the rational and efficient use of life resources, CNSA will continue to upgrade and expand. The infrastructure will be upgraded to improve the efficiency of the system and user experience. In addition, new data types associated with the omics data such as sequence, protein, metabolism, expression, annotation, clinic, image, will be gradually added to the database to enrich our database and meet the needs of more users in the future.

## Acknowledgements

We gratefully thank other colleagues in the CNGB who helped to create and maintain the CNSA.

## References

1. Samuel GN, Farsides B. (2017) The UK’s 100,000 Genomes Project: manifesting policymakers’ expectations. New genetics and society, 36, 336–53.

2. International Cancer Genome C, Hudson TJ, Anderson W, et al. (2010) International network of cancer genome projects. Nature, 464, 993–8.

3. Tomczak K, Czerwinska P, Wiznerowicz M. (2015) The Cancer Genome Atlas (TCGA): an immeasurable source of knowledge. Contemporary oncology, 19, A68–77.

4. Levy M, Chen Y, Clarke R, et al. (2020) Socioeconomic differences in health-care use and outcomes for stroke and ischaemic heart disease in China during 2009-16: a prospective cohort study of 0.5 million adults. The Lancet Global health, 8, e591–e602.

5. Exposito-Alonso M, Drost HG, Burbano HA, et al. (2019) The Earth BioGenome project: opportunities and challenges for plant genomics and conservation. The Plant journal: for cell and molecular biology.

6. Cochrane G, Karsch-Mizrachi I, Takagi T, et al. (2016) The International Nucleotide Sequence Database Collaboration. Nucleic acids research, 44, D48–50.

7. Kodama Y, Mashima J, Kosuge T, et al. (2018) DNA Data Bank of Japan: 30th anniversary. Nucleic acids research, 46, D30–D5.

8. Cook CE, Bergman MT, Cochrane G, et al. (2018) The European Bioinformatics Institute in 2017: data coordination and integration. Nucleic acids research, 46, D21–D9.

9. Sayers EW, Beck J, Brister JR, et al. (2020) Database resources of the National Center for Biotechnology Information. Nucleic acids research, 48, D9–D16.

10. Droege G, Barker K, Seberg O, et al. (2016) The Global Genome Biodiversity Network (GGBN) Data Standard specification. Database: the journal of biological databases and curation, 2016.

11. Terry SF. (2014) The global alliance for genomics & health. Genetic testing and molecular biomarkers, 18, 375–6.

12. Neumann J, Brase J. (2014) DataCite and DOI names for research data. Journal of computer-aided molecular design, 28, 1035–41.

13. Wang B, Liu F, Zhang EC, et al. (2019) [The China National GeneBank horizontal line owned by all, completed by all and shared by all]. Yi chuan = Hereditas, 41, 761–72.

14. Liu H, Wei J, Yang T, et al. (2019) Molecular digitization of a botanical garden: high-depth whole-genome sequencing of 689 vascular plant species from the Ruili Botanical Garden. GigaScience, 8.

15. Zou Y, Xue W, Luo G, et al. (2019) 1,520 reference genomes from cultivated human gut bacteria enable functional microbiome analyses. Nature biotechnology, 37, 179–85.

16. Chen S, Zhou Y, Chen Y, et al. (2018) fastp: an ultra-fast all-in-one FASTQ preprocessor. Bioinformatics, 34, i884–i90.

